# Adaptive Expression of Engrams by Retroactive Interference

**DOI:** 10.1101/2023.03.17.533126

**Authors:** Livia Autore, James D. O’Leary, Clara Ortega-de San Luis, Tomás J. Ryan

## Abstract

Long-term memories are stored as stable configurations of neuronal ensembles, termed engrams. While investigation of engram cell properties and functionality in memory recall has been extensive, less is known about how engram cells are affected by forgetting. We describe a form of interference-based forgetting using an object memory behavioral paradigm. By using activity-dependent cell labelling, we show that although retroactive interference results in decreased engram cell reactivation during recall trials, optogenetic stimulation of the labelled engram cells is sufficient to induce memory retrieval. Forgotten engrams may also be reinstated via the presentation of similar or related environmental information. Furthermore, we demonstrate that engram activity is necessary for interference to occur. Taken together, these findings indicate that retroactive interference modulates engram expression in a manner that is both reversible and updatable. Retroactive inference may constitute a form of adaptive forgetting, where in everyday life new perceptual and environmental inputs modulate the natural forgetting process.

## INTRODUCTION

Forgetting is an ubiquitous phenomenon in the animal world and can be defined as the loss of a learned behavioral response. While forgetting is commonly considered as a deficit of memory function due to its association with pathological states, an alternative emerging perspective considers forgetting as an adaptive function of the brain that may contribute to learning, and memory updating. Recent findings suggest that forgetting is an active process that involves new plasticity that modulates the functionality of specific memory traces in order to promote adaptive behavior [1–4].

The enduring physical changes occurring in neurons and synapses that represent specific information acquired during learning are termed *engrams* and constitute the physical trace of a memory [5–7]. Engram cells are operationalized as ensembles of cells that contribute to the storage of a specific memory, undergo plasticity during learning, and are likely distributed across brain regions [6, 8]. Using activity-based labelling techniques, extensive studies have described how engram cells contribute to various stages of memory formation and function including encoding, consolidation, reconsolidation, extinction and retrieval [8–11]. However, the fate of engram cells following forgetting still remains poorly understood [3, 6, 12]. Engram labelling technology allows us opportunity to investigate the nature of forgetting, for example showing that memory can still be artificially retrieved in the case of pathological conditions such as pharmacologically disrupted memory consolidation [11], Alzheimer’s [13, 14], as well as memory loss due to development and aging [14–18]. These findings are corroborated by clinical observations where previously forgotten information can be retrieved, in the example of spontaneous recovery following extinction or in periods of lucid memory recall despite Alzheimer’s pathology [19]. The survival of memory engrams supports the idea of an active form of forgetting, a process that could reversibly silence previously encoded memories [2]. However, few studies have focused on the features and characteristics of a forgotten engram, or have tested how forgetting may be modulated by direct experience.

A long-standing hypothesis is that many forms of forgetting are due to multiple memory traces encoded closely in time, competing to be consolidated in the brain [1, 20–22]. Retroactive interference constitutes an ideal model to test this hypothesis and to investigate the nature of natural forgetting because it allows experimental control of the interfering stimulus. Retroactive Interference occurs when the encoding of new specific information attenuates memory of previously encoded memories. Recognition-based behavioral tasks are well suited to study retroactive interference in rodents [23–26], because they allow us to study competing similar memories with the same emotional value[27].

Here, we developed an object-context behavioral paradigm to study interference forgetting in the mouse [23]. In this task mice are asked to recognize an object that has been displaced from its original context. We focused on the contextual engram that allows mice to discriminate the displaced and the non-displaced object. The time mice spend exploring the displaced object can be used as a measure of the contextual memory of the previous experience. We labelled a contextual engram in the hippocampus dentate gyrus (DG) of mice and studied its activation and function following interference forgetting [28–30]. The resulting findings support the hypothesis that competition between engrams mediates retroactive interference and that the forgotten memory trace can be reactivated by both natural and artificial cues as well as updated with new information. Furthermore, our data show that engram activation is necessary for forgetting to happen, consistent with the idea of forgetting as a complex and active process that requires engram cell activity.

## RESULTS

### Retroactive interference causes memory impairment and is caused by the competition of two memory engrams

We trained mice in an object-context recognition paradigm (**Figure 1a**). In this task, mice are required to learn the association between a pair of identical objects and a context and are tested 24 h later to discriminate between the learned object that has been placed in a novel context (Nov) from the Familiar one (Fam) that has not been displaced. The quantification of exploration of familiar and novel objects or conspecifics offers a within-subject control allowing us to draw robust conclusions. After two days of habituation to each of the two contexts, totalling 4 days, mice underwent a 10 min Acquisition during which they were exposed to an identical pair of objects in Context A and 1 h later an Interference phase during which they were presented with a new pair of objects in Context B. 24 h later they were tested in Context B to recognize the Nov object. A control group did not receive the Interference session and another control group was tested in Context B to test for the retroactivity of interference effect. While the levels of exploration during Acquisition and Interference phases were comparable in the three experimental groups (**Figure S1a-b**), data in **Figure 1b** show that the group that did not receive the interference session explored the novel object significantly more, while the Interference group was not able to discriminate between the two. The group tested in Context B explored the Nov object significantly more (the circle) validating the retroactive interference paradigm that we used, demonstrating that only the memory of Context A is impaired, and not of Context B (the impairment is exclusively retroactive). This result is confirmed by the Discrimination Index (**Figure 1c**).

**Figure 1.**
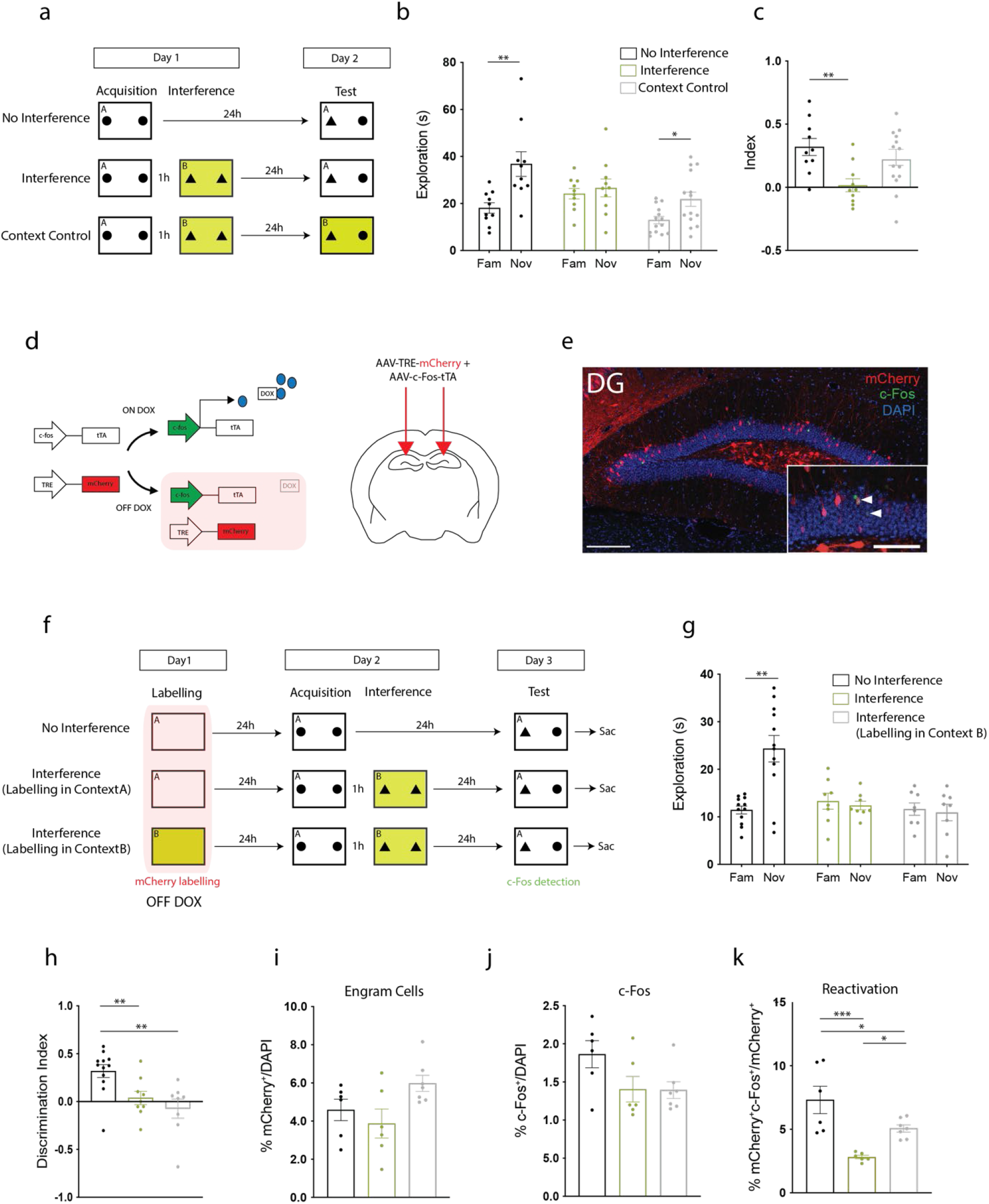
Retroactive interference causes memory impairment and drives the encoding of a new memory engram. **a.** Behavioral schedule. Acquisition and interference take place on Day 1 and mice are tested 24 h later in either the acquisition or interference context. **b.** Memory recall on Day 2. Time mice spent exploring the Familiar and the Novel object in the No Interference (N=10), Interference (N=10), Test Context B (N=14) groups. **c.** Memory index: (Novel-Familiar)/ (Novel+Familiar) for the three experimental groups. **d.** Doxycycline-dependent engram labelling strategy (left). Diagram showing the coronal section of a brain slice. The two arrows point at the site of viral injection in DG. **e.** Representative image of mCherry, c-Fos and the overlap of the two stainings in DG. DAPI in blue. White arrows: co-labelling. Scale bar: 250 μM. Scale bar in the magnified picture 125 μM. **f.** Behavioral schedule. On Day 1 mice are placed in Context A or B for labelling. 24 h later they undergo Acquisition and interference. On Day 3 Test takes place in Context A. **g.** Memory recall on Day 3. Time spent exploring the Familiar and the Novel object by No Interference mice (N=12), Interference mice (N=8) and Interference mice labelled in Context B. **h.** Memory index. Object discrimation index for 3 experimental groups. **i.** mCherry^+^ cell counts in DG. **j**. c-Fos^+^ cell counts in DG. **k**. mCherry^+^c-Fos^+^ cell counts in DG. Data presented as mean ± SEM. Unpaired t-test or one-way ANOVA Tukey; *P < 0.05, **P < 0.01, ***P < 0.001.

To test the hypothesis of interference forgetting being mediated by two competing memory traces, using a DOX-dependent labelling strategy **(Figure 1d-e)** we tagged engram cells in the DG of mice exposed to Context A and B and then trained them in the object-context recognition task (**Figure 1f**). 45 min after test subjects were perfused, we compared the level of engram reactivation in the different experimental groups. **Figure 1g** shows the behavioral performance of the three groups. As expected, the control group explored the novel object significantly more than the familiar one while the two groups that received interference were not able to discriminate between the two. This result is also confirmed by the memory index (**Figure 1h**). The number of engram cells labelled during acquisition (mCherry^+^ cells) and the number of cells activated during test (c-Fos^+^ cells) was similar in all three experimental groups (**Figure 1i-j**). However, the group that did not experience interference and the Interference group labelled in context B showed an increased percentage of reactivation of the original engram during test (**Figure 1k**), meaning that recall of the memory preferentially reactivates the original engram, whereas interference forgetting leads to the reactivation of the interference engram (Context B). These data support the idea that interference drives the encoding of a new memory engram that competes and suppresses the activation of the original memory and is retrieved during test, impairing the recall of the original trace.

### Retroactive interference forgetting can be rescued by reexposure and updated with new information

The impairment in memory retrieval that followed interference led us to test whether it was possible to rescue the behavioural performance by reexposing briefly mice to the initial object-context pair before test. Our hypothesis was that if the original memory trace persists in the DG after interference, a short reexposure (that *per se* is not enough to form a behaviourally observable memory, either 5 min or 1 h prior to test **Figure S2a-b**) might bring the neural ensemble back to an active state. At the same time, we wanted to assess how malleable a forgotten memory trace is, testing whether during this reexposure we could enrich the original memory with new information.

For this purpose, we injected a viral cocktail into mice to label engram cells for Context A with mCherry (**Figure 2a-b**) and compared the engram reactivation and memory performance of three experimental groups: a group that received interference, a group that received a brief reexposure (5 min) to the original object-context pair 5 min before test, and finally a group of mice that was reexposed for 5 min to a misleading pairing (**Figure 2c**). The behavioral performance shown in **Figure 2d** evidences that the group that received interference explored the Nov and Fam objects equally, while we were able to rescue the memory performance in the group that received reexposure to the initial object-context pair, as mice were able to distinguish them. Interestingly, the group that was reexposed to the updated object-context pairing spent more time exploring the pseudo-Fam object, confirming our hypothesis that during reexposure a memory becomes more labile and prone to updating with new information. This behavioral result is also confirmed by the memory index (**Figure 2e**). To assess if the effect of reexposure would persist over time, we also reexposed animals for 5 min to the original object-context memory 24 h before test and concluded that this does not have an effect in the long term (**Figure S2c-d**).

**Figure 2.**
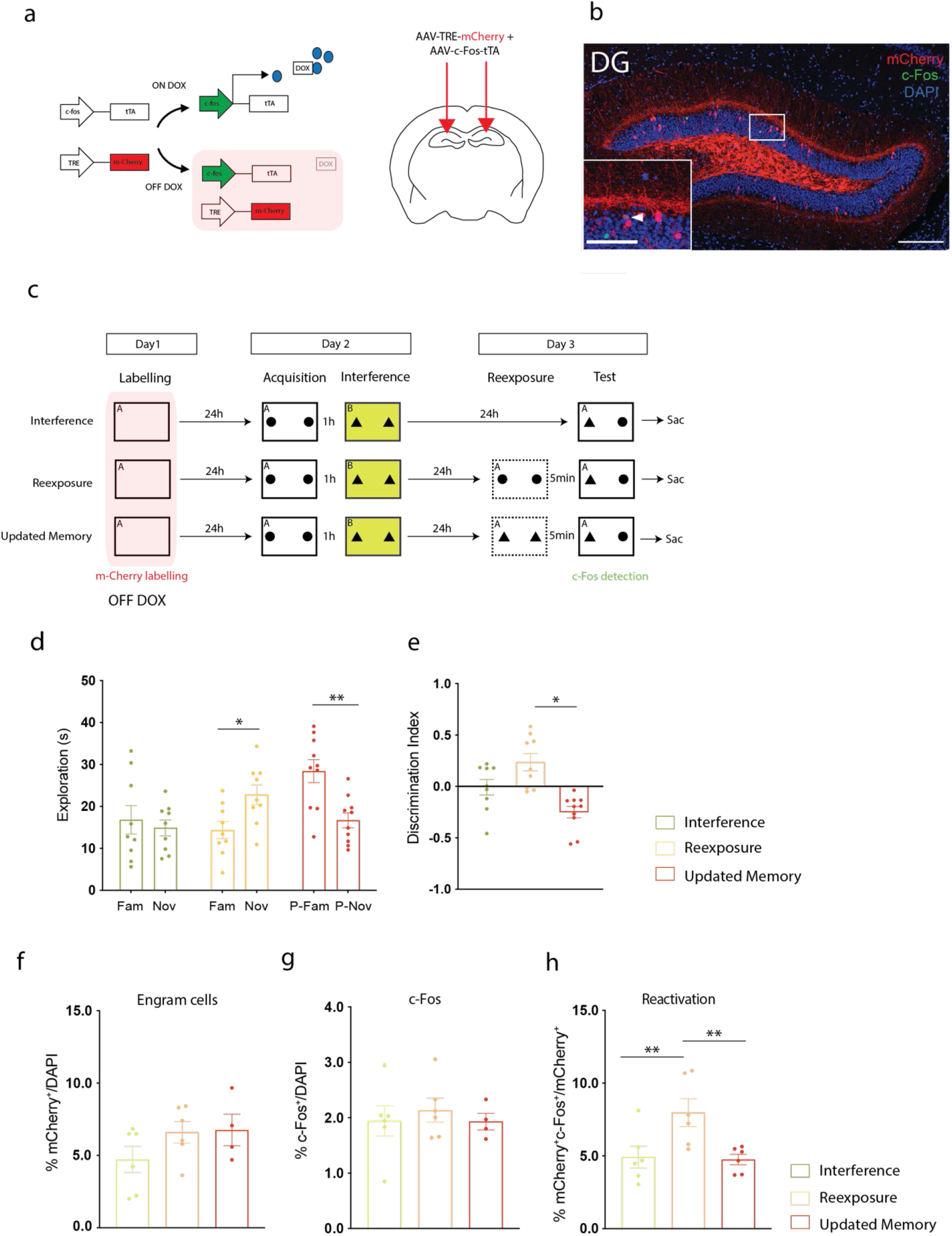
Memory loss induced by retroactive interference can be rescued by reexposure and updated with new information. **a.** Doxycycline-dependent engram labelling strategy (left). Diagram showing the coronal section of a brain slice. The two arrows point at the site of viral injection in DG. **b.** Representative image of mCherry, c-Fos and the overlap of the two stainings in DG. DAPI in blue. White arrows: co-labelling. Scale bar: 250 μM. Scale bar in the magnified picture 125 μM. **c.** Behavioral schedule. On Day 1 mice are placed in Context A or B for labelling. 24 h later they undergo Acquisition and interference. On Day 3 Test takes place in Context A without any reexposure session (Interference group), after a 5 min Reexposure session to the original object-context pair (Reexposure group) or after a 5 min Reexposure to a ‘false’ object context pair. **d.** Memory recall on Day 3. Time spent exploring the Familiar and the Novel object by Interference mice (N=9), Reexposure mice (N=9) and Update Reexposure animals (N=10). **e.** Memory index for the three experimental groups. **f.** mCherry^+^ cell counts in DG. **g.** c-Fos^+^ cell counts in DG. **h.** mCherry^+^c-Fos^+^ cell counts in DG. Data presented as mean ± SEM. Unpaired t-test or one-way ANOVA Tukey; *P < 0.05, **P < 0.01, ***P < 0.001.

The number of engram cells labelled during acquisition (mCherry^+^ cells) and the number of cells activated during test (c-Fos^+^ cells) was similar in all three experimental groups (**Figure 2f-g**). In accordance with their memory performance, the group that received reexposure to the original object-context pair showed a higher level of reactivation of engram cells compared to the other two experimental groups in support of the hypothesis that reexposure restores accessibility to memory engrams. Even though reexposure to a false object-context pair induced higher exploration of the pseudo-Fam than the pseudo-Nov object, the engram reactivation detected after test was comparable to the interference group, suggesting that reexposure to the misleading cues might lead to the recruitment of different engram cells (**Figure 2h**).

### Artificial reactivation of engram cells in DG rescues interference-induced forgetting

Activity of memory engrams in DG has been shown to be necessary and sufficient for memory expression [8, 31]. Hence, we tested whether we could restore memory performance in mice following interference by optogenetically reactivating the contextual engram. To this aim, we labelled engram cells using a cocktail of AAV9-cfos-tTA + AAV9-TRE-ChR2 and implanted optic fibers in the DG of subjects[32] (**Figure 3a-b**). Context A engram cells were labelled during the OFF DOX period and mice were later exposed to acquisition and interference. 24 h later, 5 min before test, they were placed in a neutral context for 5 min in the presence of the initial pair of objects and engram cells were stimulated in DG through light stimulation. To test for unspecific effects of the surgery and fibers implantation a group of mice was reexposed to the same conditions without light being delivered. To test whether reexposure to both components (object + context) of the memory is necessary, we also reexposed subjects before test to only objects or only context and assessed their memory (**Figure 3c**). What stands out in **Figure 3d** is that memory performance in the opto-stimulated group is rescued and comparable to the reexposure group, with mice exploring significantly more the novel object than the familiar one, indicating that engrams in DG encode the contextual component of the object-context memory. Animals reexposed to only objects or only context as well as the No Light control mice were not able to discriminate between novel and familiar objects, since they explore equally the two categories. These results are confirmed by the memory index shown in **Figure 3e**.

**Figure 3.**
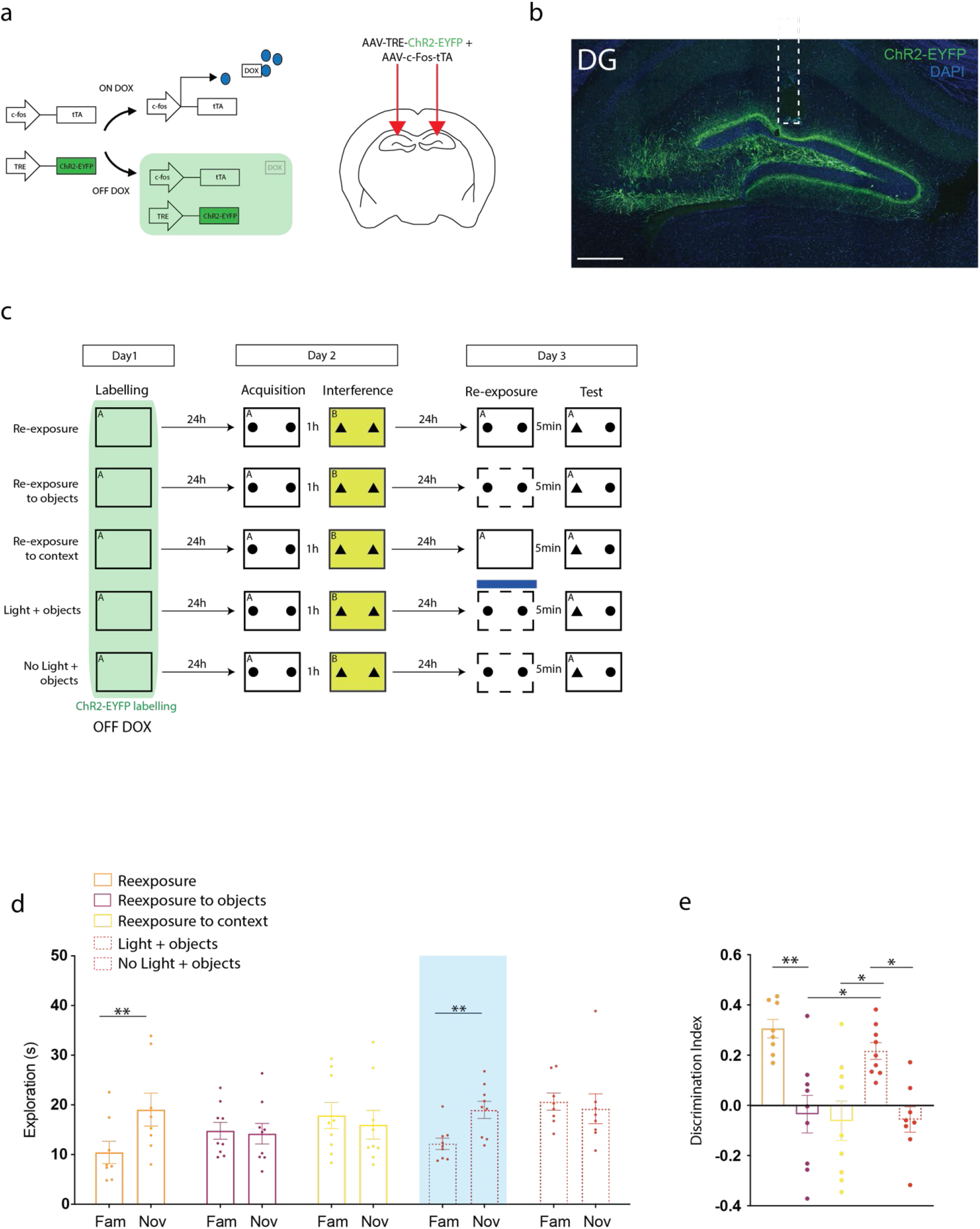
Optogenetic reactivation of a contextual engram before test rescues mice memory performance. **a.** Doxycycline-dependent engram labelling strategy (left). Diagram showing the coronal section of a brain slice. The two arrows point at the site of viral injection in DG. **b.** Representative image of a coronal slice of the DG with ChR2-EYFP labelling (green) and DAPI (blue). White dashed rectangle signalling implant location. Scale bar: 250 uM **c.** Behavioral schedule. On Day 1 mice are placed in Context A for labelling. 24 h later they undergo Acquisition and interference. On Day 3 Test, prior to test, subjects are reexposed to the original object+context pair/only objects/only context/object+opto-stimulation/object+no light depending on the experimental group they are assigned to. 5 min later, test takes place in Context A. The blue thick line indicates the opto-stimulation session. **d.** Memory recall on Day 3. Time spent exploring the Familiar and the Novel object by the Reexposure mice (N=8), Reexposure to only objects (N=9), Rexposure to only context (N=9), Reexposure to objects + light stimulation (N=9), and Reexposure to objects + no light(N=8). **e.** Memory index for the different experimental groups. Data presented as mean ± SEM. Unpaired t-test or one-way ANOVA Tukey; *P < 0.05, **P < 0.01, ***P < 0.001.

### Interference forgetting is an active process that requires the activation of the forgotten engram

The evidences reported so far showed that engram cells are still present in the DG following interference and can be altered by experience. This might suggest that forgetting is an active process of selection and inhibition of memory traces to be consolidated, inhibited, or updated. Because of this, we asked whether the activity of the original object-context memory engram is necessary during interference for forgetting to happen. Mice have been injected with a viral cocktail (AAV9-cfos-tTA + AAV9-TRE-ArchT-GFP) to label engram cells encoding for Context A with an inhibitory opsin (**Figure 4a-b**) and implanted with optic fibers in DG. Following labelling, they were trained and exposed to the interference stimuli. Light-induced inhibition of engram cells was performed throughout the duration of the 10 min interference session to block the activation of Context A engram cells during this period. A control group of mice received the same treatment except for the light delivery that was not performed (**Figure 4c**). 24 h later, subjects were tested and their memory performance was assessed. Interestingly, light-induced inhibition of Context A engram cells during interference prevented forgetting as it’s shown by the significant increase in exploration of the novel object compared to the familiar one in the Light group. The same effect cannot be observed in mice in the No Light group who explored the two objects equally (**Figure 4d**). All these data are confirmed by the memory index (**Figure 4e**).

**Figure 4.**
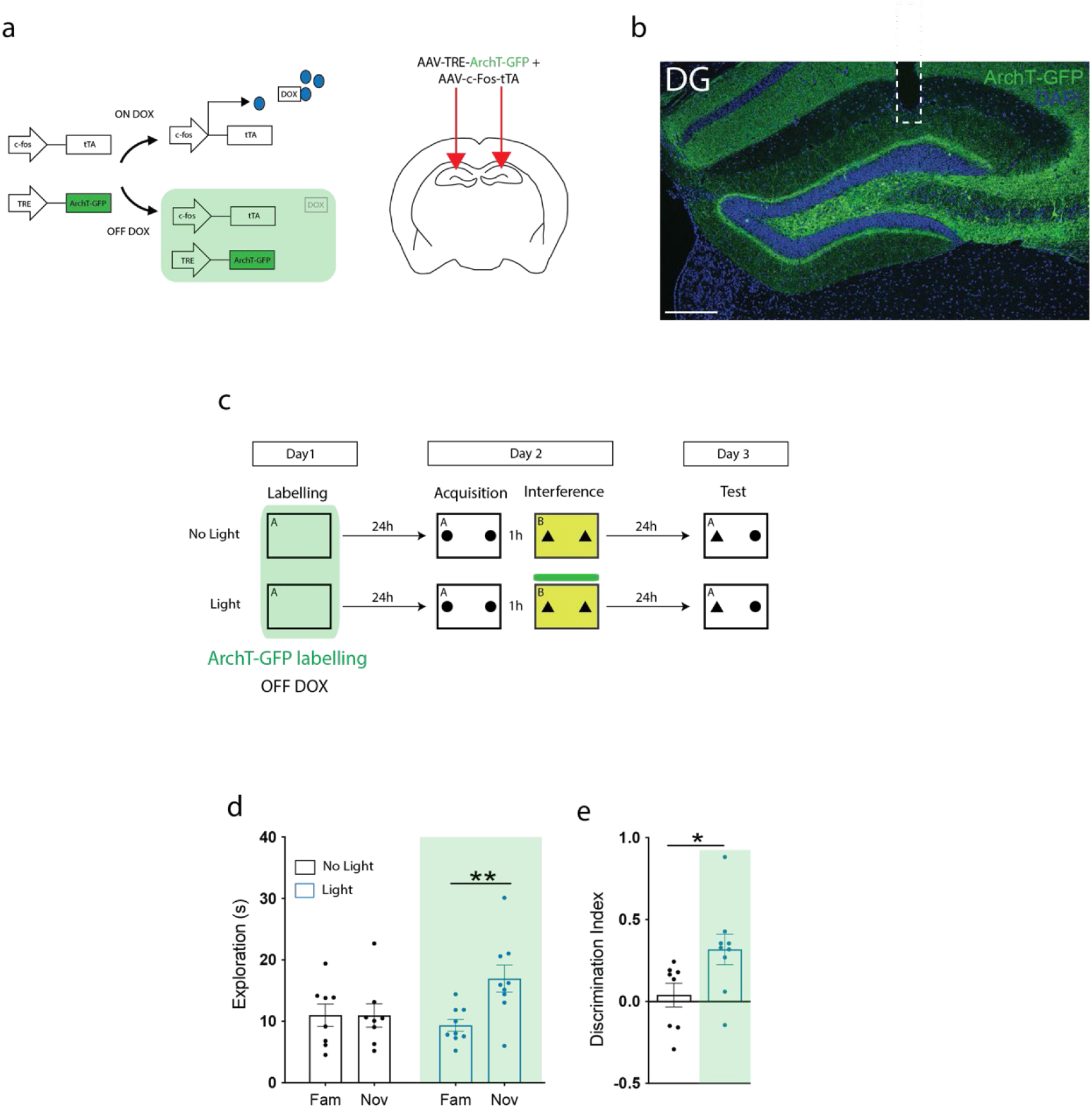
Optogenetic inhibition of a contextual engram during interference prevents forgetting. **a.** Doxycycline-dependent engram labelling strategy (left). Diagram showing the coronal section of a brain slice. The two arrows point at the site of viral injection in DG. **b.** Representative image of a coronal slice of the DG with ArchT-GFP labelling (green) and DAPI (blue). White dashed rectangle signalling implant location. Scale bar: 250 uM **c.** Behavioral schedule. On Day 1 mice are placed in Context A for labelling. 24 h later they undergo Acquisition and receive opto-inhibition of the contextual memory engram during Interference. On Day 3, test takes place in Context A. The yellow thick line indicates the opto-inhibition session. **d.** Memory recall on Day 3. Time spent exploring the Familiar and the Novel object by the No-light control mice (N=8) and the Light group (N=9). **e.** Memory index for the two experimental groups. Data presented as mean ± SEM. Unpaired t-test or one-way ANOVA Tukey; *P < 0.05, **P < 0.01, ***P < 0.001.

### Exposure to an enriched environment causes interference in a contextual fear conditioning paradigm

To test whether our findings were replicable when the two competing memories are distinct with different emotional value, we developed another interference protocol utilizing a contextual fear conditioning paradigm. We labelled engram cells with mCherry (**Figure 5a-b**) when mice were exposed to Context A on Day 1. Then, 24 h later mice received three footshocks in Context A. The control group was then placed back in their homecage, while the interference group was immediately placed for 2 h in an enriched environment cage with their cagemates. 24 h later all mice were placed back in Context A and their freezing levels were quantified as a measure of memory expression (**Figure 5c**). As an effect of interference, mice who were exposed to the enriched environment immediately after training showed a significant reduction in freezing levels during test, compared to the control group. While the number of engram cells labelled during acquisition (mCherry^+^ cells) and the number of cells activated during test (c-Fos^+^ cells) was similar in the two experimental groups (**Figure 5e-f**), mice in the interference group showed a decrease percentage of reactivation of the original engram during test (**Figure 5g**).

**Figure 5.**
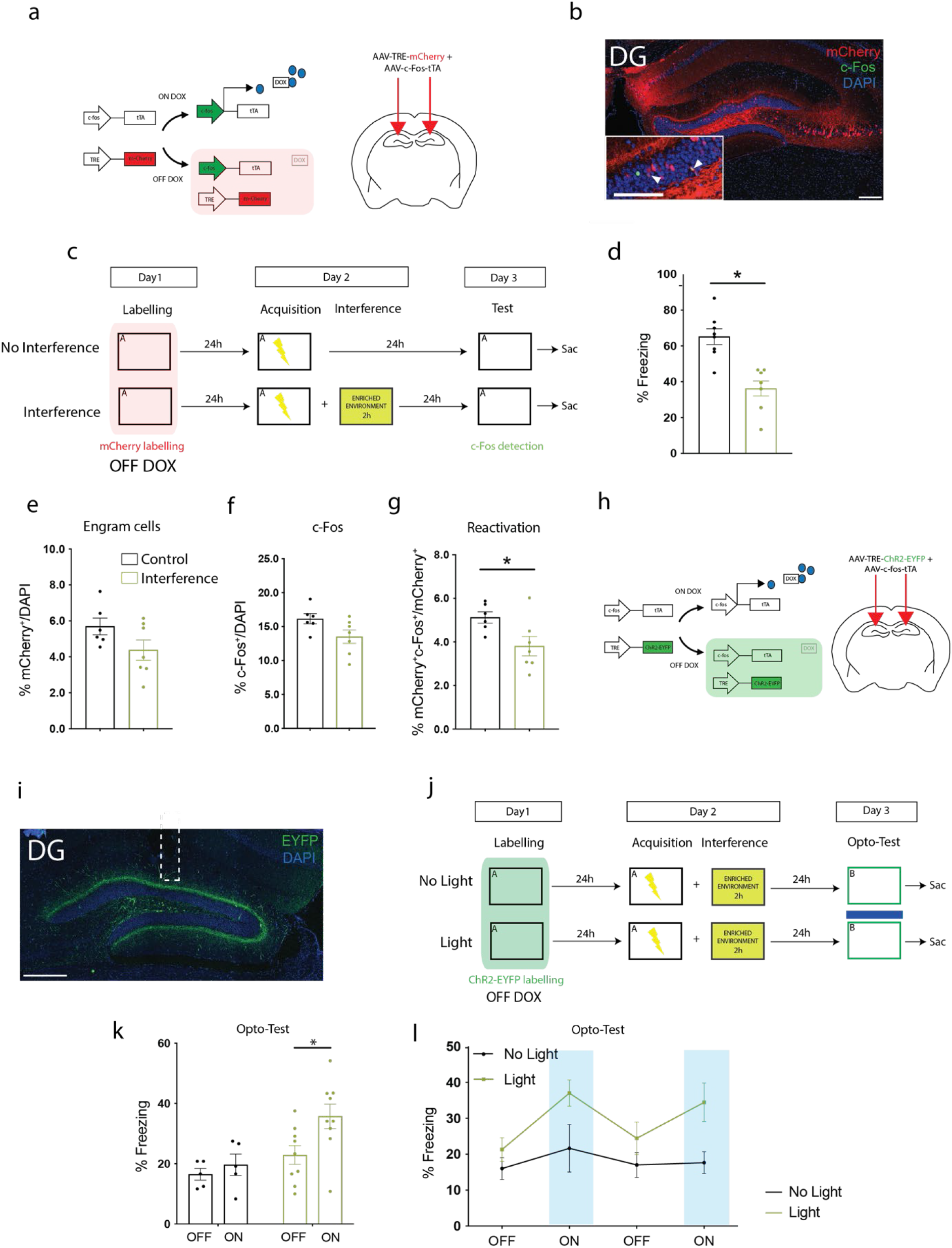
Exposure to an enriched environment causes interference forgetting of a contextual fear memory. **a.** Doxycycline-dependent engram labelling strategy (left). Diagram showing the coronal section of a brain slice. The two arrows point at the site of viral injection in DG. Behavioral schedule. **b.** Representative image of mCherry, c-Fos and the overlap of the two stainings in DG. DAPI in blue. White arrows: co-labelling. Scale bar: 250 μM. Scale bar in the magnified picture: 125μM **c.** Behavioral schedule. On Day 1 mice are exposed to Context A for 3 min for engram labelling. 24 h later, they’re placed back in Context A. On Day 2 they are trained in Context A receiving a footshock. Interference mice are placed in an enriched environment for 2 h immediately after training. Animals are tested in Context A for 3 min on Day 3. **d.** Memory Recall test on Day 3. Percentage of mice spent freezing by the Control group (N=8) and the Interference group (N=7). **e.** mCherry^+^ cell counts in DG. **f.** c-Fos^+^ cell counts in DG. **g.** mCherry^+^c-Fos^+^ cell counts in DG. **h.** Doxycycline-dependent engram labelling strategy (left). Diagram showing the coronal section of a brain slice. The two arrows point at the site of viral injection in DG. **i.** Representative image of a coronal slice of the DG with ChR2-EYFP labelling (green) and DAPI (blue). White dashed rectangle signaling implant location. **j.** Behavioral schedule for opto-stimulation in DG. On Day 3 mice are tested in Context B receiving opto-stimulation. **K-l.** Memory Recall test on Day 3. Percentage of time mice spent freezing by the No Light group (N=5) and the Light group (N=9) in the Neutral Context B. Data presented as mean ± SEM. Unpaired t-test or one-way ANOVA Tukey; *P < 0.05, **P < 0.01, ***P < 0.001.

To artificially reactivate the forgotten memory, we then injected a cocktail of AAV9-cfos-tTA + AAV9-TRE-ChR2 into mice and implanted optic fibers in the DG of mice (**Figure 5h-i**). After labelling Context A engram cells, mice were fear conditioned and placed in the enriched environment for interference. 24 h later they were tested in a neutral Context B in a 12 min test session during which they received four 3 min-epochs of blue light on and off (**Figure j**). The No Light group was tethered to the laser but did not receive any light stimulation. As illustrated by **Figure 5k-l**, mice in the Light group froze significantly more during the ON epochs, supporting the hypothesis that the artificial stimulation of the forgotten memory is sufficient to elicit fear expression.

Furthermore, no difference in light-induced freezing was observed between an Interference group and a No Interference group (Figure S5). We injected (**Figure S3a-b**) and trained two additional groups of mice that underwent fear conditioning. One group was used as a No Interference control and the other was placed into the interference cage for 2 h. On Day 3 mice were tested for Natural Recall and on Day 4 engram cells were artificially stimulated in Context B (**Figure S3c**). **Figure S3d-f** show freezing levels of the two experimental groups during natural recall and opto-test, respectively.

## DISCUSSION

Our results demonstrate that retroactive interference represents a form of natural forgetting, that is an adaptive and active process caused by experience, and is due to the competition of multiple memory engrams that become more or less accessible according to environmental changes. Through the use of engram labelling strategies and optogenetics we characterized memory engrams following retroactive interference and showed that *forgotten* memories persist in the DG even when memory performance is impaired. Previous studies have already provided evidences of engram survival and memory rescue by direct optogenetic stimulation for different types of amnesia [7, 11, 13–18] but here, for the first time, this was investigated for retroactive interference forgetting. Our data fit well the existing literature supporting the critical role of the hippocampus and the DG in the encoding and retrieval of contextual information [28]. Particularly, protein synthesis in the DG has been shown to be necessary for object-context long term memory formation [29].

Interference is possibly one of the most common types of forgetting in our everyday life as we are continuously exposed to a constant flux of information that necessarily leads to the competition of multiple memory traces with different values and features [3]. Our data provide evidence that when retroactive interference occurs in an object-context recognition task, the memory engram encoding the forgotten trace is retrievable by natural and artificial reexposure to cues, suggesting that during reexposure new plasticity mechanisms might be responsible for the strengthening of the original memory expression. This hypothesis is in accordance with previous research showing that engram reactivation during reexposure to cues following learning is necessary for memory expression [8, 11, 33, 34]. Interestingly we found that following interference, an engram can change if subjects are reexposed to misleading cues, inducing the consolidation of a ‘false’ or updated memory. The idea that ‘poorly-encoded’ engrams are preserved in a labile state that can be altered in multiple directions is supported by experimental and modelled data showing that in a fear conditioning paradigm overgeneralized memories are not crystallized, and can be modified during recall by environmental circumstances [35, 36]. We showed that the level of reactivation of the original engram during the recall of the updated memory was decreased, suggesting that the behavioral reversal that we observed in mice is reflected by an update in the composition of neurons taking part to the memory engram. This evidence suggests that forgotten memory traces are still labile and can be subjected to new plasticity.

The hypothesis that forgetting is an active process, characterized by plasticity mechanisms similar to the ones occurring upon learning, is supported by our data showing that during interference the activity of an engram is necessary for its own forgetting. One explanation could be that engrams cells have to activate some intrinsic molecular pathways to signal that they need to be silenced or become inaccessible [3]. For example, it has been shown that in *Drosophila*, inhibition of the small G-protein Rac1 in neurons slows down memory decay from 2 hr up to 24 hr [37] and in mice the level of Rac1 within engram cells affects memory maintenance until two weeks after memory formation [38].

A second option could be that the competing engram (Context B) is partially recruiting neurons from the first ensemble during memory allocation. Thus, those neurons need to be active to allow the formation of a functional and prevailing competing engram. This hypothesis is supported by data showing that the overlap between two memory traces is influenced by the time interval separating the two experiences: the closer in time the two events are encoded, the more the potential for overlap [39, 40]. Even though our overlap study shows that following interference there is a reduction in the activation of the original memory engram, we could still observe a small degree of overlap between neurons activated in the acquisition context and test (2.66% reactivation). It might be plausible that when these shared neurons are inhibited during interference, the correct formation of an interference engram is prevented and thus the consequent forgetting.

As previously mentioned, the object-context task that we developed is ideal to investigate the biological substrates of interference forgetting as it allows us to look at memories closely-encoded in time that are also similar in content and emotional value, and gives us temporal control of natural forgetting. However, our hypothesis that forgetting is mediated by competition of memory traces is supported by the fact that we were able to replicate our results in a contextual fear conditioning task. We developed a novel fear conditioning-based retroactive interference paradigm that utilizes an enriched environment to interfere with the contextual experience. We showed that not only memory interference occurs in the case of fear memory with a strong emotional value, but also that we were able to artificially retrieve it, confirming the presence of the original memory engram in the DG. These fear conditioning results constitute a strong validation of our forgetting model.

An important study by Poll and colleagues [14] showed that in a mouse model of Alzheimer Disease the impairment of hippocampus-dependent memories is due to the activation of a new ensemble (‘novelty-like engram cells’) that is recruited when mice are not able to recall the original memory and causes pathological forgetting. These findings are in accordance with our data as the experience of the second object-context pair that mice receive in our paradigm represent a novel ensemble that causes the memory impairment. It is tempting to speculate that pathological forgetting in cases such as Alzheimer Disease is due to the aberrant activation of natural forgetting processes such as seen in retroactive interference.

In conclusion, here we proposed a model of interference forgetting based on the competition of multiple distinct ensembles in DG where engram cells that encode for a forgotten memory are activated to become more or less accessible depending on environmental cues. We speculate that one of the causes that might determine the retrievability of an engram is the prediction/error ratio. A developing perspective suggests that new learning proceeds as a function of this ratio [41, 42]. When an event is mispredicted mice will form a new, separate, neural representation of that item. The greater is the misprediction, the more is the differentiation between the two representations. In our retroactive interference paradigm, the presentation of a similar but different object-context pair represents an expectancy violation that contributes to the formation of a new engram and promotes active forgetting. According to our findings we suggest that interference constitutes a robust model for forgetting processes that is generalizable to different forms of memory. Furthermore, we showed that engram activation is necessary for forgetting to occur, demonstrating that neuronal activity and plasticity are crucial for this process. Future research is needed to elucidate what molecular mechanisms are involved in these processes and if and how they can be modulated to determine the degree and rate of forgetting.

Forgetting (and remembering) can be considered as a gradient of engram expression [3] ranging from severely reduced memory performance, such as in Alzheimer Disease [13, 14], to abnormal memory abundance, potentially as in Autism Spectrum Disorder [18]. Retroactive interference may lie at the centre of this spectrum as it constitutes a reversible form of natural forgetting that allows memory expression to be changed as a function of everyday experience.

The continuous flow of environmental changes leads to the encoding of multiple engrams that compete for their consolidation and expression. Some may persist undisturbed, some will be subjected to interference by new incoming and prevailing information. However, the interfered memories can still be reactivated by surrounding cues leading to memory expression or by misleading or novel experiences ending up in an updated behavioural outcome.

Conceiving interference as a model of reversible memory suppression provides a key for human studies to elucidate and interpret the neurobiological basis of multiple types of active forgetting [43–45].

## ACKNOWLEDGMENTS

We thank Tamara Boto, Bianca Silva, and Rasmus Bruckner for useful discussions. This work was funded by the Irish Research Council, Science Foundation Ireland, and the European Research Council.

## AUTHOR CONTRIBUTIONS

L.A., J.OL., and T.J.R. conceived the scientific design. L.A. conducted the experiments and analysed the data. L.A., J.OL., C.O.S. and T.J.R. interpreted the results. L.A. and T.J.R. wrote the original draft. L.A., C.O.S., J.OL. and T.J.R. reviewed and edited the manuscript.

## SUPPLEMENTAL FIGURE TITLES AND LEGENDS

**Figure S1.**
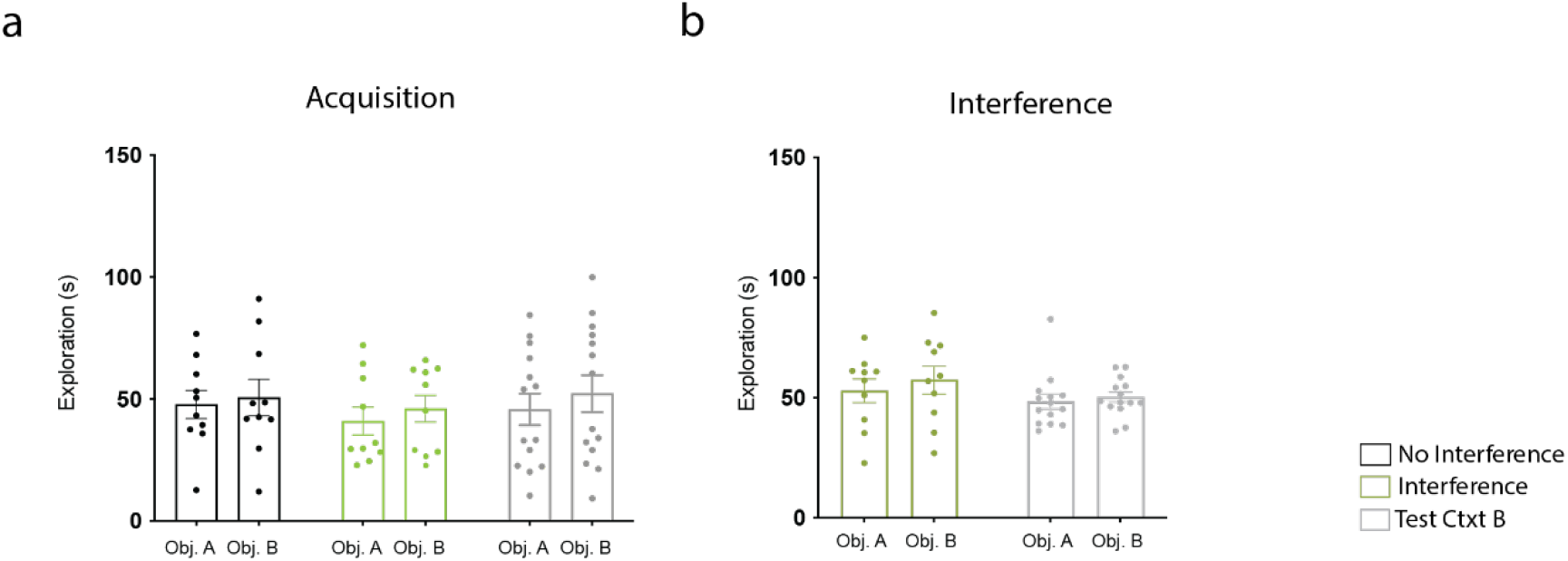
During acquisition and interference there is no difference in the exploration of the two objects. **a-b.** Time spent by mice exploring the two objects during Acquisition and Interference.

**Figure S2.**
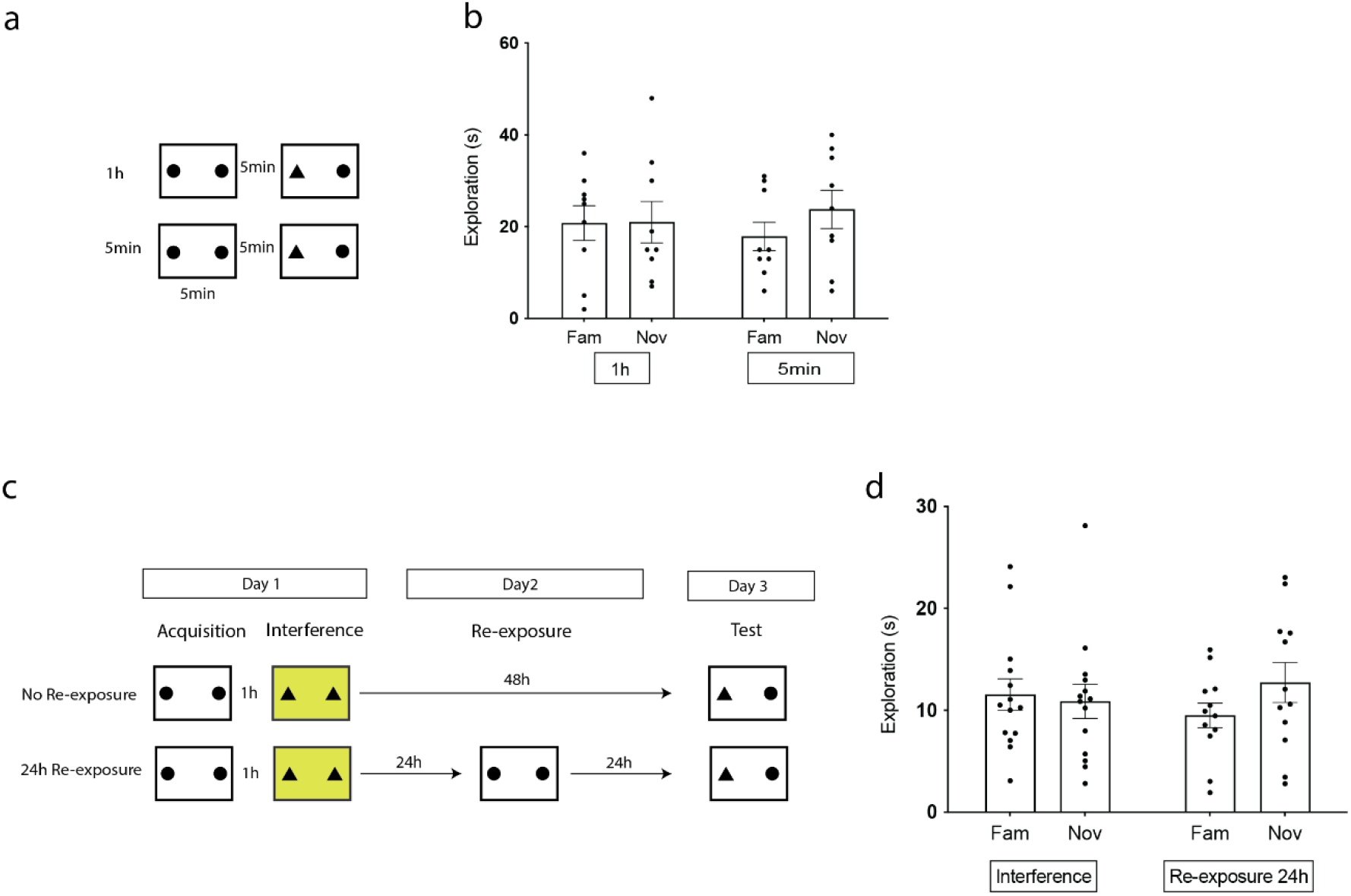
Control experiments for the effects of reexposure. **a.** Behavioral schedule. On the same day mice were exposed for 5 min (dotted arena indicates brief experience) to context A with an identical pair of objects and tested for novel object recognition 1 h and 5 min later. **b.** Memory recall data. There was no difference in the exploration of the two categories of objects in either group. **c.** Behavioral schedule. On Day 1 mice undergo Acquisition and interference. On Day 2 the Reexposure group is reexposed to the first object-context pair for 5 min, the No Reexposure group does not receive any reexposure. On Day 3 mice are tested for 5 min in Context A. **d.** Time spent by the mice exploring the two categories of object. There was no difference in the exploration of the novel and the familiar object in either group. Data presented as mean ± SEM. Unpaired t-test or one-way ANOVA Tukey; *P < 0.05, **P < 0.01, ***P < 0.001.

**Figure S3.**
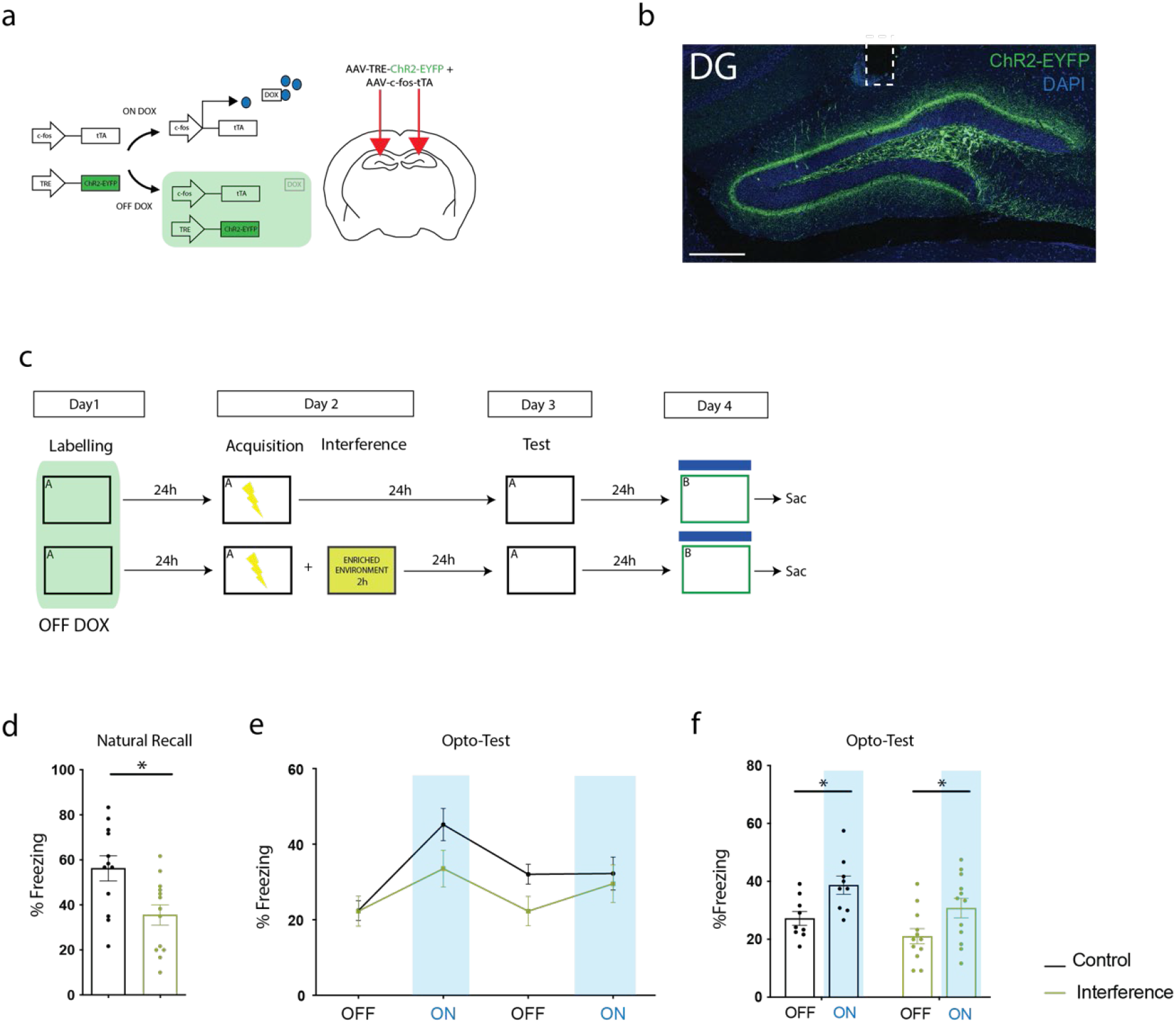
Comparison between No Interference and Interference groups during natural recall and opto-test. **a.** Doxycycline-dependent engram labelling strategy (left). Diagram showing the coronal section of a brain slice. The two arrows point at the site of viral injection in DG. **b.** Representative image of a coronal slice of the DG with ChR2-EYFP labelling (green) and DAPI (blue). White dashed rectangle signalling implant location. **c.** Behavioral schedule for opto-stimulation in DG. On Day 3 mice are tested in Context A for natural recall. On Day 4 mice are tested in Context B receiving opto-stimulation. **d.** Memory Recall test on Day 3. Percentage of time mice spent freezing by the No Interference (N=12) and Interference group (N=12). **e-f.** Memory Recall opto-test on Day 4. Percentage of time mice spent freezing by the two groups during the ON and OFF epochs. Data presented as mean ± SEM. Unpaired t-test or one-way ANOVA Tukey; *P < 0.05, **P < 0.01, ***P < 0.001.

## MATERIAL AND METHODS

### Subjects

C57Bl6/J mice have been used for these experiments. They were all males between the age of 7 and 12 weeks. Animals were housed in groups of five in cages with a tunnel, food and water *ad libitum*. The animal room was kept at a constant temperature of 22°C. Mice were exposed to a 12 h light/dark cycle and each step of the experiments was conducted during the light phase. All the procedures related to mouse care and behavioural experiments were carried out in accordance with Health Products Regulatory Authority (HPRA) Ireland guidelines.

### Engram labelling strategy and stereotactic surgery

The labelling strategy used for these experiments is DOX-dependent. Mice were taken OFF DOX 36-40 h before the labelling session. Following labelling, DOX food was immediately reintroduced into their diet. The viral cocktails injected for each experiment are summarised in the following table:

| EXPERIMENT | VIRUSES INJECTED | TOTAL VOLUME<br>INJECTED | TITER |
| --- | --- | --- | --- |
| Figure 1, 2, 5 | AAV9-c-fos-tTA (1:10)<br>AAV9-TRE-mCherry (6:10) | 300 nL x hemisphere | 4.72x10<sup>13</sup>gc/mL<br>5.00x10<sup>13</sup>gc/mL |
| Figure 3 | AAV9-c-fos-tTA (1:10)<br>AAV9-TRE-ChR2 (4:10) | 300 nL x hemisphere | 4.72x10<sup>13</sup>gc/mL<br>2.74x10<sup>13</sup>gc/mL |
| Figure 4 | AAV9-c-fos-tTA (1:10)<br>AAV9-TRE-ArchT (4:10) | 300 nL x hemisphere | 4.72x10<sup>13</sup>gc/mL<br>6.69x10<sup>13</sup>gc/mL |

Stereotactic surgery was performed 8-14 days before starting the behavioural procedures. Mice were anaesthetized with 500 mg kg^-1^ avertin and head fixed on the stereotactic. Each animal underwent bilateral craniotomy and two holes were drilled at −2.00 mm anteroposterior (AP), ± 1.35 mm mediolateral (ML) for DG injections.

The viral mix was injected using a microsyringe pump (UMP3; WPI) and a 10 μL Hamilton syringe (701LT; Hamilton) at −2.00 mm dorsoventral (DV) coordinate. Injection speed was 60 nL/min. When needed, a bilateral fiber implant (Doric) was fixed to the stereotaxic adapter and was lowered in the brain at −1.85 mm and secured with dental cement and a protective cap. Mice were allowed to recover for 8 days before any behavioural procedure was performed.

### Immunohistochemistry

Mice were injected intraperitoneally with an overdose of Sodium Pentobarbital (50 μL) and perfused transcardially with PBS and then with 4% paraformaldehyde in PBS (Sigma Aldrich). After 24 h incubation in paraformaldehyde 4%, brains were then sliced using a vibratome. Five coronal slices per mouse were selected from the structure of interest.

For the immunostaining, slices were permeabilized three times for 10 min in PBS. Then, they were incubated in a blocking solution (PBT 0.2%, NGS 10%) for 1 h at room temperature. Successively, they were incubated with primary antibody (in PBT0,2% +NGS 3%) overnight at 4°C while shaking. On the following day, slices underwent three 10 min washes in PBT 0.1% and then incubated with secondary antibody (PBT 0.2%, NGS 3%) for 2 h in the dark at room temperature. Then, after three additional 10 min washes in PBT 0.1%, slices were stained with DAPI (1:1000 in PBS) for 15 min at room temperature. Slices were finally mounted on superfrost slides using Vectashield-DAPI (Vector Laboratories).

The antibodies used for the c-Fos staining were rabbit anti-c-Fos (Synaptic Systems – Ref. Number: 226003) (1:500) and anti-rabbit Alexa-488 (Invitrogen - Ref. Number: A11034) (1:500). ArchT-EYFP and ChR2-EYFP were stained using a primary chicken anti-GFP IgY fraction (Invitrogen – Ref. Number: A10262) and then a rabbit anti-chicken Alexa-488 IgG (Invitrogen – Ref. Number: A11039) (1:500).

### Implants verification and cell counting

Slices were mounted onto super frost slides using Vectashield DAPI. Implant locations for each animal were visualized and verified using an Olympus BX51 upright microscope. When the implant was not targeted correctly into the region of interest (DG) the animal was excluded from the behavioral analysis.

To measure the reactivation of engram cells during test, sections (5 x mouse) were imaged on a Leica Confocal at 20x magnification. For each image, the total number of cells in the DG was calculated (DG area/cell area). Then, the number of engram cells (mCherry^+^ cells) and c-Fos+ cells was counted. These were divided by the total number of DAPI-positive cells to estimate the percentage of c-Fos^+^ cells and mCherry^+^ cells in the DG. To quantify the reactivation of engram cells, the number of cells positive for both mCherry and c-Fos was divided by the number of mCherry^+^ cells. The reactivation value for each mouse was calculated by averaging the reactivation for each slice.

### Behavioral procedures

#### Object-context Memory task

Object-context task took place in 2 different contexts. Context A was a rectangular 40×20×30 cm arena with a grey floor and white walls decorated with black triangles. Context B was an arena of the same size but with black and white striped walls. The objects used for this task were a purple bottle (height: 15 cm, diameter 5 cm) and a small white, blue and red garden gnome (height: 12 cm). The two objects were always presented in pairs (of identical or different objects) and each of them was placed facing the wall on one of the two short sides of the arena. All objects were counterbalanced within each experiment.

Mice were habituated to each context for 10 min for two consecutive days before the behavioural procedure started.

During acquisition, mice were placed in context A in the presence of two identical objects. 1 h later, interference took place in context B in the presence of another pair of identical objects. On test day, mice were placed back in context A and allowed to explore 2 different objects: the one they had previously seen in context A and one that they’d previously seen in context B. Object exploration was used as a measure of the animals’ memory performance and was defined as the time the subject’s nose was < 2 cm from the object.

The arena was cleaned with trigene (Distel Laboratory Disinfectant) between one mouse and the following.

In the presence of a reexposure session, mice were placed in Context A together with the same objects they had seen in acquisition or interference context, respectively for reexposure and updated memory groups.

#### Optogenetic engram stimulation

The optogenetic stimulation session took place in a neutral context with white walls and white floor in the presence of two identical objects. The optical fibre implant was connected to a 450 nm diode fibre light source (Doric LDFLS 450/080). The stimulation protocol consisted of constant stimulation throughout the whole session with a frequency of 20 Hz and a pulse width of 15 ms.

#### Optogenetic engram inhibition

took place during the interference session. Mice performed the task while being connected to a 520 nm diode fibre light source for inhibition. The duration of the session was 10 min and during the whole duration of the session the mouse was allowed to explore the arena and light was delivered continuously.

#### Contextual Fear Conditioning

The Contextual Fear Conditioning arena consisted in a 31 x 24 x 21 cm chamber located into a soundproof box. The chamber had a removable grid floor and grey walls, except for one transparent wall and black opaque triangular perspex ceiling. The context was sprayed with 1% acetic acid (Sigma-Aldrich) diluted in tap water used as olfactory cue. 24 h before training, subjects were exposed for 3 min to the conditioning contexts and were free to explore the arena to allow the labelling of engram cells encoding for the context. Then, they were returned to their homecage and DOX food was reintroduced in their diet. Training day took place in the same arena 24 h later. The training protocol had a total duration of 300 s. After 3 minutes of habituation, mice were presented with 3 shocks (0.2s, 0.75mA) separated by 1 min interval (at 180, 210 and 270s respectively).

On the test day mice were reexposed to context A for 180 s and their freezing behaviour was assessed. No shock was delivered during the test.

#### Optogenetic Recall Test

The opto-stimulation session took place in a different context characterised by white floor, white circular walls and benzaldehyde 0.25% in ethanol (Sigma-Aldrich) as an olfactory cue. Subjects were placed in the arena and the optical fibre implant was connected to a 450 nm diode fibre light source (Doric LDFLS 450/080). The duration of the session was 12 min and during the whole duration of the session the mouse was allowed to explore the arena. The stimulation protocol consisted in 2 light-off and 2 light-on epochs: during the first 3 minutes of the session and from min 6 to 9 the laser was off. From min 3-6 and 9-12 the laser was switched on and mice received the light stimulation with a frequency of 20 Hz and a pulse width of 15 ms.

#### Interference by Enrichment

To interfere with the contextual learning experience, mice were placed in a large cage for 2 h with their cagemates. The cage was enriched with bedding, 2 nests, paper, two red tubes, lego bricks, a jar, a running wheel and food. After 2 h subjects were placed back in their homecage.

### Analysis and statistics

All the experiments were analysed in blind and videos randomised before manual scoring. Statistics were conducted using Prism 9.00 (Graphpad software). Unpaired Student T-test was used for comparison between groups and one-way ANOVA followed by a Tukey post hoc test was used for multiple comparisons.

